# Seeking rhythmic patterns in microglial cells within the circadian pineal gland from male rats

**DOI:** 10.1101/2025.01.10.632375

**Authors:** Carlos L. Freites, Martin Avila, Juan B. Amiotti, Estela M. Muñoz

**Affiliations:** Laboratory of Basic and Translational Neurobiology, Institute of Histology and Embryology of Mendoza (IHEM), National University of Cuyo (UNCuyo), National Scientific and Technical Research Council (CONICET), Mendoza, Argentina

**Keywords:** pineal gland, microglia, microglial heterogeneity, IBA1-positive cells, daily rhythms

## Abstract

Microglia, the innate immune cells of the brain, constitute a highly dynamic cell population that displays several functions influenced by the light:dark (L:D) cycle. Within the pineal gland (PG), a key organ of the circadian timing system, microglia actively participate in its development and homeostasis. However, little is known about their rhythmic features in this circadian organ. This study aimed to elucidate morphological and functional phenotypes of pineal microglial cells at two time points of the L:D cycle. We performed immunofluorescence staining and confocal microscopy on paraffin-embedded pineal sections from 3– and 18-month-old Wistar rats, analyzing samples collected at midday (ZT6) and midnight (ZT18). Our results showed that the density and spatial distribution of IBA1^+^ microglial cells did not vary between ZT6 and ZT18 in the 3-month-old PG. However, these cells exhibited reduced size and amoeboid-like shapes, along with a reduction in the expression of the phagocytosis inhibitor SIRP alpha, at ZT18 compared to ZT6. Moreover, IBA1^+^ cells were immunoreactive for the lysosomal marker CD68 and the autophagic marker LC3B at both ZTs. Additionally, contacts with PAX6^+^ cells were detected in the two ZTs analyzed. Interestingly, IBA1^+^ cells showed attenuated or abolished rhythmicity in morphological parameters and SIRP alpha expression levels in the 18-month-old PG. Our findings suggest that microglia undergo transitions into alerted states at night, characterized by small, rounded shapes and enhanced phagocytic capacity. This adaptation likely prepares them to respond to potential invading agents and other insults in the PG, which could impact the nocturnal melatonin production.

**Key points:**

- Microglia within the rat pineal gland exhibit daily morphological and functional adaptations, enhancing their effectivity to respond against potential challenges in a time-dependent manner.
- Oscillatory features of pineal microglia diminish with aging, suggesting an attenuation of their functions.
- Nocturnal reactivity of microglia within the pineal organ may influence its physiology, potentially affecting melatonin synthesis.

## 1. Introduction

The capacity to sense time enables organisms to initiate molecular, cellular, and physiological changes that facilitate adaptation and anticipation to fluctuating environmental conditions between day and night. All these changes are mediated by endogenous timing mechanisms distributed throughout various cells and tissues, known as biological clocks (Gliech & Holland, 2020; Starnes & Jones, 2023). The circadian timing system (CTS) ensures that a variety of biological processes are synchronized and constrained within a time frame that cycles approximately every 24 h, known as circadian rhythms (Reppert & Weaver, 2002; Schulz & Steimer, 2009).

In mammals, a complex and multisynaptic circuit known as photoneuroendocrine system transduces information related to the photoperiod into a hormonal output, the synthesis and release of melatonin (MEL) by the pineal gland (PG) (Korf et al., 1998; Maronde & Stehle, 2007). The hormone MEL is an indoleamine rhythmically produced at night by the pinealocytes (Farias Altamirano et al., 2019), which are the predominant cell population of the PG. Its production is tightly controlled by the central clock located within the suprachiasmatic nuclei of the hypothalamus (Borjigin et al., 2012; Hastings et al., 2019). Sympathetic nerves emerging from the superior cervical ganglia innervate the PG, releasing norepinephrine at nighttime, which stimulates MEL synthesis (Avila et al., 2023; Schomerus & Korf, 2005). Once released into the periphery, MEL acts as a molecular signal that synchronizes multiple biological events (Pandi-Perumal et al., 2006; Pevet & Challet, 2011).

As the primary immune cells of the central nervous system (CNS), microglial cells directly participate in the development and homeostasis of the PG (Muñoz, 2018, 2022). We have previously demonstrated that IBA1^+^ microglia regulate PAX6^+^ precursor-like cell density, and remodel blood vessels and nerve fibers in the developing rat PG (Ibañez Rodriguez et al., 2016). Moreover, pineal microglial cells are also highly sensitive to microenvironmental fluctuations, as reflected in their ability to react efficiently and differentially to surgical and pharmacological stimuli via changes in number and phagocytic capacity (Ibañez Rodriguez et al., 2018).

Recent evidence based on single-cell RNA sequencing analysis has identified two microglial subtypes, termed α– and β-microglia, within the rat PG. Notably, α-microglia display a small subset of transcripts that are upregulated at daytime compared to the β-microglia subpopulation. Despite this, the transcriptomic profile of pineal microglial cells shows minimal regulation by the light:dark (L:D) cycle. This contrasts with pinealocytes, which exhibit highly rhythmic gene expression (Mays et al., 2018). Nevertheless, microglia throughout other brain regions, such as cortex and hippocampus, can display oscillatory functions, particularly those related to interactions with neurons, phagocytosis, and cytokine release (Choudhury et al., 2020; Fonken et al., 2015; Hayashi et al., 2013; Takayama et al., 2016).

While it is known that microglia exhibit oscillatory functions, their daily dynamics in a circadian organ such as the PG are far from conclusive. In the present study, we applied a multiple immunofluorescence labeling protocol followed by three-dimensional confocal imaging to characterize microglial phenotypes influenced by the L:D cycle in the young adult and aged rat PG. Our data suggest that pineal microglia are multifaceted cells with rhythmic features, and these daily adaptations might contribute to the proper functioning of the organ.

## 2. Materials and Methods

### 2.1 Animals

Young adult (3-month-old; P3m, where P indicates postnatal age) and aged (18-month-old; P18m) male Wistar rats (*Rattus norvegicus*; RRID:RGD_13508588) were used in this study. Animals were bred and housed in a temperature– and humidity-controlled room at the local facilities of the Institute of Histology and Embryology of Mendoza. They were maintained on a 12:12 h L:D cycle and provided *ad libitum* access to food and water. The turning on of the room lights was established at 7 AM and was considered as ZT0 (ZT, *Zeitgeber* time). The euthanasia was performed either at ZT6 (middle of the day) or ZT18 (middle of the night). In the case of sacrifices at ZT18, they were conducted in a darkroom illuminated with low-intensity red light as performed before (Castro et al., 2015; Farias Altamirano et al., 2023). All experimental procedures were implemented according to the National Institutes of Health’s Guide for Care and Use of Laboratory Animals, the Animal Research: Reporting *In Vivo* Experiments (ARRIVE), and were approved by the Institutional Animal Care and Use Committee at the School of Medicine, National University of Cuyo, Mendoza, Argentina (Protocol IDs 74/2016, 152/2019, and 218/2022). All efforts were made to minimize the number and suffering of the animals during all experimental procedures.

### 2.2 Tissue preparation and immunohistochemistry

Rats were euthanized by decapitation after being CO_2_-anesthetized, and the PG of each animal was immediately collected either at ZT6 or ZT18. Individual PGs were processed as described before (Castro et al., 2015; Farias Altamirano et al., 2023; Ibañez Rodriguez et al., 2016). Briefly, PGs were fixed by immersion in 4 % (w/v) paraformaldehyde (PFA) in 1X phosphate-buffered saline (PBS) at 4°C for 24 h. After fixation, the glands were rinsed in 1X PBS and dehydrated in increasing concentrations of ethanol solutions. PGs were finally processed and embedded in yellow modified paraffin wax (Histoplast; Biopack, Buenos Aires, Argentina, Cat# CAS 8002-74-2). 10 μm-thick sections were cut from tissue blocks at random orientations from the central area of each PG using a Microm HM 325 microtome (Thermo Fisher Scientific Inc., Waltham, USA; RRID:SCR_020259).

For immunofluorescence staining, slide-mounted tissue sections were deparaffinized, hydrated, and boiled in 0.01 M sodium citrate buffer (pH 6) containing 0.05 % (v/v) Tween-20 for 30 min for antigen retrieval. Non-specific labeling was avoided by using a blocking solution [3 % (w/v) bovine serum albumin (BSA) and 1 % (v/v) Triton X-100 in 1X PBS] for 1 h at room temperature (RT) in a humid chamber. Immunolabeling was performed by incubating the tissue slices in an antibody solution [3 % (w/v) BSA and 1 % (v/v) Triton X-100 in 1X PBS] containing primary antibodies (Table 1), overnight at 4°C in a humid chamber. After incubation with the primary antibodies, sections were rinsed in 1X PBS and then incubated in the antibody solution containing secondary antibodies (Table 2) at RT for 2 h. Finally, slices were rinsed in 1X PBS and then coverslipped with N-propyl-gallate mounting medium [2 % (w/v) N-propyl-gallate (Sigma-Aldrich, St. Louis, USA, Cat# P3130), and 90 % (v/v) glycerol in PBS] containing 0.15 % (w/v) propidium iodide (PI; Sigma-Aldrich, Cat# P4170) for cell nuclear staining. Alternatively, nuclear counterstaining was carried out with 4’,6-diamidino-2-phenylindole dihydrochloride (DAPI; Thermo Fisher Scientific Inc., Cat# D1306) in a dilution of 1:600. Staining controls were routinely performed by the omission of primary antibody incubation. Images were captured using an Olympus FV1000 confocal microscope (Olympus America Inc., Center Valley, USA; RRID:SCR_020337), and processed with the open-source Fiji software (https://fiji.sc/; RRID:SCR_002285).

**Table 1:**
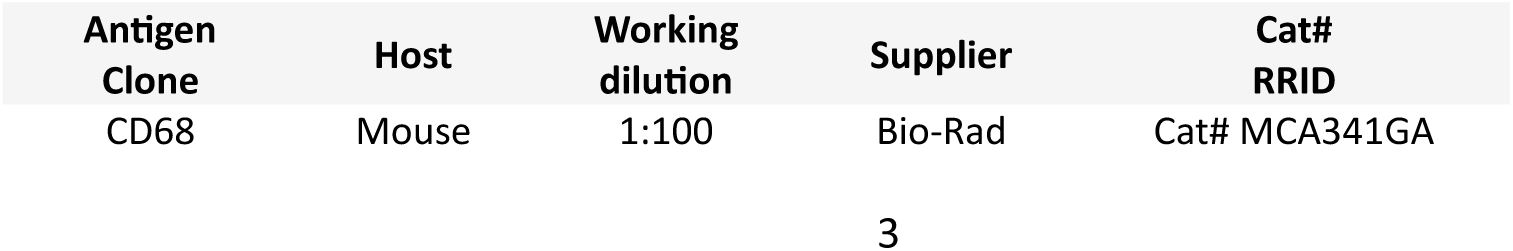
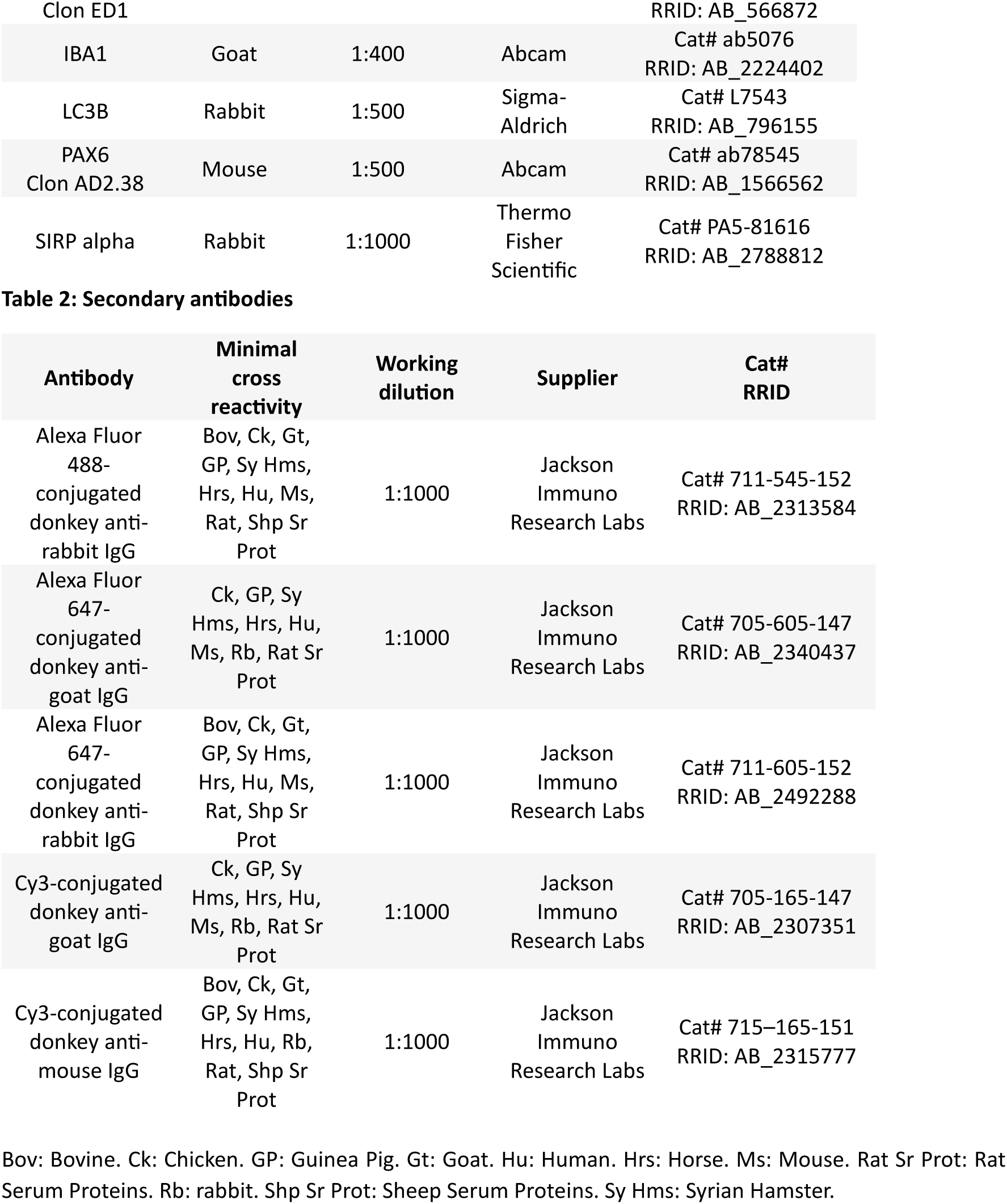
Primary antibodies.

### 2.3 Antibody characterization

The anti-CD68 monoclonal antibody generated in mice (Bio-Rad, clone ED1, Cat# MCA341GA) was raised against the rat CD68 (cluster of differentiation 68) protein derived from spleen cells. The antibody ED1 recognized a heavily glycosylated protein of ∼90-110 kDa expressed in monocytes and macrophages (Dijkstra et al., 1985). ED1 antigen is located mostly in cytoplasmic granules such as phagosomes, and on the cell surface (Damoiseaux et al., 1994). We have previously shown that the anti-CD68 antibody is suitable for targeting microglial cells within the rat PG (Ibañez Rodriguez et al., 2016, 2018). Other studies have demonstrated its expression in hippocampus and spinal cord (Cope et al., 2016; Park et al., 2024). The IBA1 (ionized calcium-binding adapter molecule 1) protein was identified using an anti-IBA1 polyclonal antibody (Abcam, Cat# ab5076) generated in goat and raised against a synthetic peptide corresponding to the C-terminus of the human protein (amino acids 135-147). The antibody specifically recognized a single band at ∼17 kDa on Western blots of rat brain lysates (see the manufacturer’s datasheet). The anti-IBA1 antibody has been extensively used for identifying microglial cells across various CNS regions (Guneykaya et al., 2018; Hu et al., 2021), including the PG, as demonstrated in our previous works (Farias Altamirano et al., 2023; Ibañez Rodriguez et al., 2016, 2018).

The anti-LC3B monoclonal antibody (Sigma-Aldrich, Cat# L7543) was generated in rabbit against a KLH-conjugated linear peptide corresponding to amino acids 2-15 from the N-terminal region of the human LC3B (microtubule-associated protein 1A/1B light chain 3B). This antibody detected a band at ∼15 kDa on Western blots of lysates from HeLa cells treated with chloroquine, and a cytoplasmic punctate staining on paraffin-embedded sections of rat and human cerebral cortex (see the manufacturer’s datasheet). The LC3B autophagic protein has been immunodetected in the rodent brain (Li et al., 2023; Vidyadhara et al., 2023). The immunostaining pattern shown herein matched that described in a previous study performed in the rat PG (Ge et al., 2021).

The PAX6 (paired box 6) transcription factor was recognized using a mouse anti-PAX6 monoclonal antibody (Abcam, clone AD2.38, Cat# ab78545). This antibody was produced by immunizing animals with a recombinant full-length protein corresponding to the human PAX6. The specificity of the antibody was verified by Western blot using protein extracts from SO-Rb50 and Y79 human retinoblastoma cell lines (Meng et al., 2014). The nuclear expression pattern of PAX6 detailed herein matched that observed in previous works on the rat PG (Ibañez Rodriguez et al., 2016, 2018), and in other studies on the cortex and retina (Cunningham et al., 2013; Gong et al., 2023).

The rabbit anti-SIRP alpha polyclonal antibody (Thermo Fisher Scientific, Cat# PA5-81616) was raised against the recombinant rat SIRP alpha (signal-regulatory protein alpha). The antibody has been tested by Western blot on protein lysates from the NIH/3T3 fibroblast cell line and by immunohistochemistry on paraffin-embedded rat tissue sections (see the manufacturer’s datasheet). The transmembrane receptor SIRP alpha has been identified via immunolabeling with other antibodies in microglia and neurons within the retina and thalamus (Jiang et al., 2022; Lehrman et al., 2018). Our immunostaining revealed a punctate pattern in pineal tissue consistent with that observed in the neuronal regions mentioned above.

### 2.4 Quantification of the number of individualized IBA1^+^ cells

Gland sections from P3m rats were immunolabeled for IBA1 and nuclear counterstained. One section per animal (N=5 glands per ZT) was stained. Imaging was performed with the Olympus FV-1000 confocal microscope, using a 40X objective lens (UPLFLN, NA: 1.30, oil immersion objective). Images were obtained at a resolution of 640×640 pixels and a scanning speed of 8 μs/pixel. Stacks of 10 optical planes (Z-step=1 μm) were projected in the Z axis for the analysis. One area of 0.05 mm^2^ was selected within each image and IBA1^+^ cells were quantified manually using the Cell Counter plugin of Fiji. The number of areas per PG section depended on the size of the gland and ranged from 10 to 14. Each area contained at least one IBA1^+^ cell body. Excluded from the counting were any IBA1^+^ signal (e.g., cytoplasmic branches), where cell nuclei could not be identified, and any IBA1^+^ cell clusters where cells could not be properly individualized.

### 2.5 Quantification of the number of IBA1^+^ cell clusters

The images used to quantify the number of individualized IBA1^+^ cells were also employed for calculating the number of IBA1^+^ cell clusters. Cluster evaluation was carried out by adapting a published protocol (Paasila et al., 2020). Clusters were counted in an area of 0.1 mm^2^. A cell cluster was defined as a group of at least three IBA1^+^ cell somas displayed within a 1600 μm^2^ area of interest. Additionally, it was required that the cytoplasm and/or branches of these cells were in proximity along the X and Y axes, as well as along the Z axis. For large-size clusters, quantification was done by digitally dividing them into multiple 1600 μm^2^ areas of interest, each considered as an individual cluster.

### 2.6 Morphometry of IBA1^+^ cells

Morphological characterization was performed on images taken from gland sections derived of P3m and P18m rats stained for IBA1 and nuclear counterstained. One section per animal (N=5 glands per ZT and per age) was stained. Images were acquired using a 60X objective lens (Plan Apo, NA: 1.42, oil immersion objective) with a 3.8x digital zoom. Each IBA1^+^ cell was completely imaged along the Z axis, and the morphometry was performed on maximum intensity Z-stack projection of the IBA1 channel. Cell clusters and cells located at the edges of the images whose branches were truncated, were not subjected to analysis. Approximately 5 to 7 IBA1^+^ cells were selected from the lower, middle, and upper regions of each tissue section, resulting in more than 50 cells per ZT and per age. Image processing was performed following a modified version of a previously established protocol by using Fiji (Young & Morrison, 2018). Briefly, IBA1-channel images were systematically subjected to Unsharp Mask, Subtract Background and Gaussian Blur Filter commands, to sharpen IBA1 signal and reduce the background. Next, images were empirically thresholded to generate binary masks. Pixels of interest corresponding to each binary mask of the former IBA1^+^ cell were selected, and morphological parameters were calculated using the same method as previously applied (Fernández-Arjona et al., 2017). The cell size (e.g., cell area and cell perimeter) and shape descriptors (e.g., circularity, solidity, and elongation index) were evaluated.

### 2.7 Analysis of the expression pattern of CD68 in IBA1^+^ cells

Gland sections (N=5 per ZT) from P3m rats co-stained for IBA1, CD68, and DAPI as nuclear marker, were imaged using a 60X objective lens with a 3.8x digital zoom. The background was subtracted from the channels corresponding to the IBA1 (Cy3-conjugated secondary antibody) and CD68 (Alexa Fluor 647-conjugated secondary antibody) markers in images generated from Z-stack projections, by using Fiji. Around 5 to 7 IBA1^+^ cells were selected from the lower, middle, and upper regions of each tissue section, thereby analyzing more than 50 cells per ZT. IBA1^+^ cells were classified according to the immunoreactivity for CD68, as IBA1^+^ CD68^-^ (no detectable levels of CD68 or levels below the detection limit based on the image acquisition settings used), or IBA1^+^ CD68^+^ (punctate pattern positive for CD68 distributed in the cytoplasm and/or branches).

### 2.8 Measurement of mean fluorescence intensity of SIRP alpha signal

Gland sections (N=4 per ZT and age) from P3m and P18m rats were immunolabeled for IBA1 and SIRP alpha, and counterstained with DAPI. Sections were imaged using a 60X objective lens with a 3.8x digital zoom. They were excited with the 405 nm, 559 nm, and 635 nm lasers to detect cell nuclei (DAPI), IBA1 (Cy3-conjugated secondary antibody), and SIRP alpha (Alexa Fluor 647-conjugated secondary antibody), respectively, with the laser power set up at 10 %. The scanning settings remained constant across all samples. Image processing and fluorescence intensity quantification were conducted by using Fiji, as instructed by an adaptation of a published protocol (Shihan et al., 2021). Stacks (10 optical planes; Z-step=1 μm) were processed as previously described to generate binary masks of the IBA1 signal. These binary masks were used to accurately define the SIRP alpha fluorescent signal in each IBA1^+^ cell. Once the perimeter of each IBA1^+^ cell was defined, this selection was transferred to the image corresponding to the SIRP alpha signal. More than 50 cells per ZT and per age were analyzed. The mean fluorescence intensity (MFI) of SIRP alpha signal was calculated as:

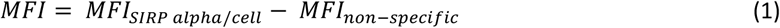

where MFI_SIRP alpha/cell_ represents the mean brightness values of SIRP alpha signal in each IBA1^+^ cell, and MFI_non-specific_ corresponds to the fluorescence intensity due to non-specific binding of the Alexa Fluor 647-conjugated secondary antibody in the absence of the primary antibody.

### 2.9 Analysis of interactions between PAX6^+^ cells and IBA1^+^ cells

Sections of PGs (N=5 per ZT) from P3m rats immunolabeled for IBA1 and PAX6, and counterstained with DAPI, were imaged using a 40X objective lens. One area of 0.05 mm^2^ was selected in each image for measurements. The number of images per gland depended on the size of each tissue section, ranging from 10 to 14. The quantification of contact events between these cell populations was performed following our established protocol (Ibañez Rodriguez et al., 2016, 2018). Contact events were defined as instances of proximity between PAX6^+^ cells and the cytoplasmic branches of IBA1^+^ cells. Additionally, IBA1^+^ amoeboid-like cells with their soma protruding towards Pax6^+^ cells were also considered. Cell-cell contacts were confirmed by analyzing optical planes individually within each Z-stack. IBA1^+^ clusters, where cells were not properly individualized, were excluded from this analysis.

### 2.10 Statistical analysis

Data were expressed as mean ± standard error of mean (SEM). The statistical analysis was customized to the type of data under examination. Data normality was evaluated using the Shapiro-Wilk test. For data that exhibited normal distribution, Student’s t tests were implemented. For data that did not show normal distribution, Mann–Whitney U tests were performed. A p<0.05 was considered significant. Analyses were conducted on four or five PGs per ZT and per age. Analyses and graphical representations were conducted using GraphPad Prism 7.00 software (http://www.graphpad.com/; RRID:SCR_002798).

## 3. Results

To study the morphological and functional changes on pineal microglial cells between two time points of the L:D cycle, we immunostained PG sections from P3m and P18m Wistar rats, sacrificed at ZT6 (midday) and ZT18 (midnight), for the classic microglia/macrophage marker ionized calcium-binding adapter molecule (IBA1) (Jurga et al., 2020). Although we successfully identified microglial cells within the PG, we do not exclude the possibility that CNS-associated macrophages were also detected (Mildenberger et al., 2022). Henceforth, we use ‘IBA1^+^ cells’ to name all phagocyte populations immunoreactive for this marker within the pineal organ.

### 3.1 Density and spatial arrangement of IBA1^+^ cells in the young adult PG

As previously reported by our group in the adult PG at ZT6 (Ibañez Rodriguez et al., 2016), we found that IBA1^+^ cells were also spread randomly throughout the P3m gland at ZT18, with no apparent influence of the L:D cycle on their abundance and distribution (Figure 1a^1^-a^3^ and 1b^1^-b^3^). Our quantitative analysis showed no significant daily differences in the number of individualized IBA1^+^ cells (11.86 ± 0.88 cells/0.05 mm^2^ and 11.41 ± 0.84 cells/0.05 mm^2^ for PGs collected at ZT6 and ZT18, respectively; p=0.4690) (Figure 1c^1^). Furthermore, pineal IBA1^+^ cells were sometimes aggregated in clusters (Figure 1a^3^ and 1b^3^). Even though these clusters did vary in size and shape, we found no significant difference in their numbers between both ZTs (0.4048 ± 0.14488 clusters/0.1 mm^2^ and 0.3590 ± 0.1855 clusters/0.1 mm^2^ for samples collected at ZT6 and ZT18, respectively; p=0.2609) (Figure 1c^2^).

**Figure 1.**
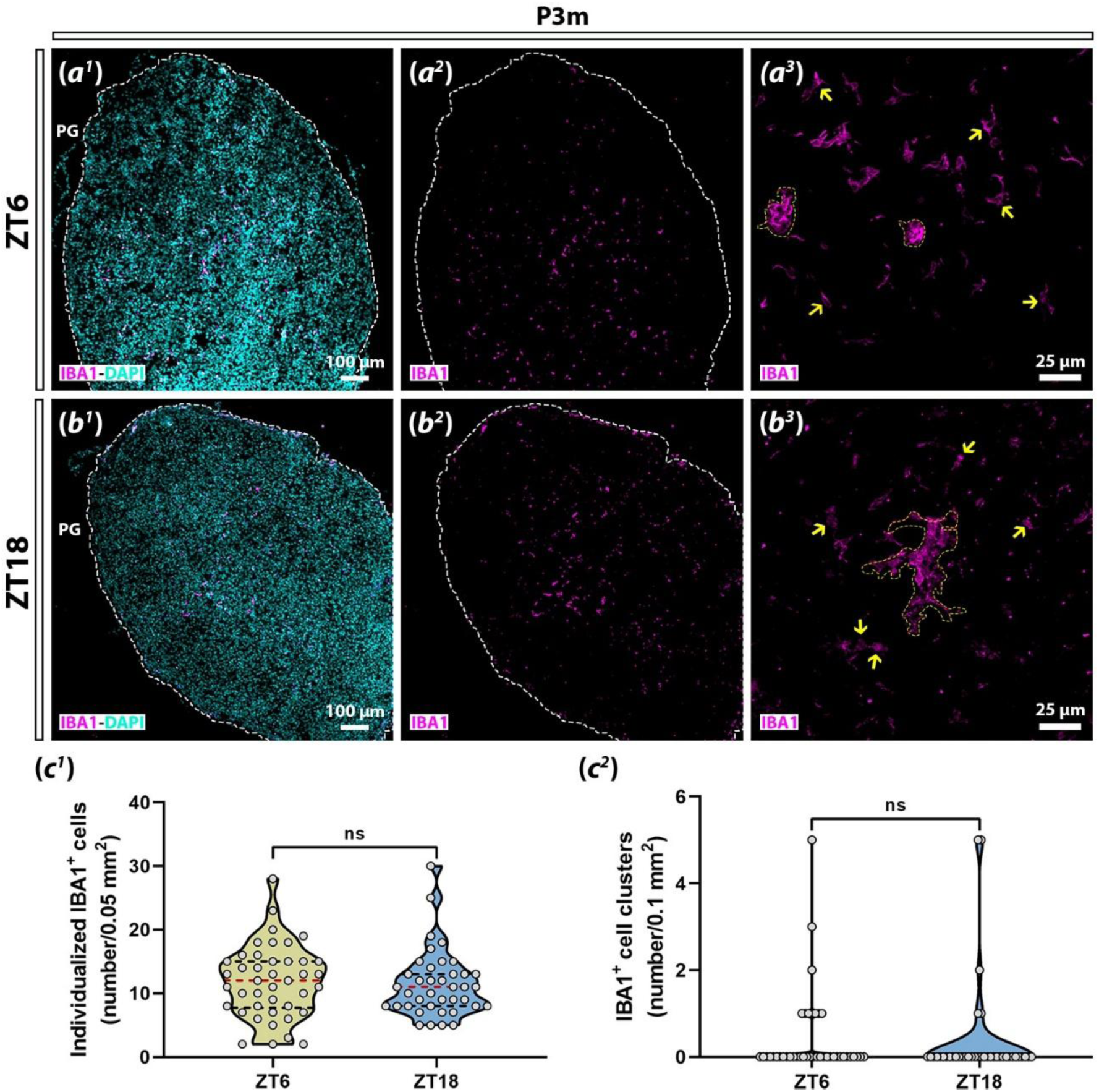
IBA1^+^ cells within the P3m PG. **(a-b)** Representative images of gland sections immunolabeled for IBA1 (magenta). DAPI (cyan) was used as nuclear marker. (a^1^, a^2^, and b^1^, b^2^) Images taken at 10X. The perimeter of each gland is delimited by a white dashed line. (a^3^ and b^3^) Images at 40X. Individualized IBA1^+^ cells are indicated by yellow arrows and IBA1^+^ cell clusters are bounded by yellow dashed lines. **(c)** Quantification of IBA1^+^ cell density. (c^1^) Number of individualized IBA1^+^ cells (Mann-Whitney U test; p=0.4690). (c^2^) Number of IBA1^+^ cell clusters (Mann-Whitney U test; p=0.2609). Data were expressed as mean ± SEM. The red dashed line in violin plots indicates the median, while the black dashed lines indicate the quartiles. ns: not significant. PG: pineal gland. P3m: 3-month-old.

### 3.2 Morphological features of pineal IBA1^+^ cells

Morphological plasticity of microglial cells has been extensively studied in a plethora of contexts. Changes in the shape of these cells often serve as reliable indicators of alterations in both their microenvironment and their functional responses (Savage et al., 2019; Vidal-Itriago et al., 2022). Considering this dynamism, we quantified morphological parameters in IBA1^+^ cells within the P3m PG, at both ZT6 and ZT18 (Figure 2a). Concerning cell size, both cell area (94.80 ± 4.97 μm^2^ and 63.59 ± 4.87 μm^2^ for glands harvested at ZT6 and ZT18, respectively; p<0.0001) and cell perimeter (110.20 ± 7.03 μm and 75.53 ± 6.14 μm; p<0.0002) exhibited significant decreases at ZT18 compared to ZT6 (Figure 2b and 2c). On the other hand, shape descriptors, such as circularity (0.17 ± 0.02 and 0.22 ± 0.02; p<0.0060) and solidity (0.54 ± 0.02 and 0.63 ± 0.02; p<0.0065), showed significantly higher values at midnight (Figure 2d and 2e). However, elongation index (2.30 ± 0.10 and 2.68 ± 0.20; p=0.5692) did not display any significant difference between both ZTs (Figure 2f). Traditionally, microglia have been morphologically classified as either ramified or amoeboid according to their functional status (Paolicelli et al., 2022). However, these morphological phenotypes—morphotypes— represent only the extremes of a broad and continuous spectrum of shapes, which are even seen to vary across different parts of the brain (Tan et al., 2020; Vidal-Itriago et al., 2022). Although microglial cells have been described as exhibiting reduced ramification within CVOs (Ibañez Rodriguez et al., 2016, 2018; Muñoz, 2022; Takagi et al., 2019), and we further demonstrated herein that IBA1^+^ cells adopt more amoeboid-like shapes in the P3m PG at midnight, we cannot overlook the presence of numerous intermediate morphotypes under normal conditions (Suppl. Figure 1).

**Figure 2.**
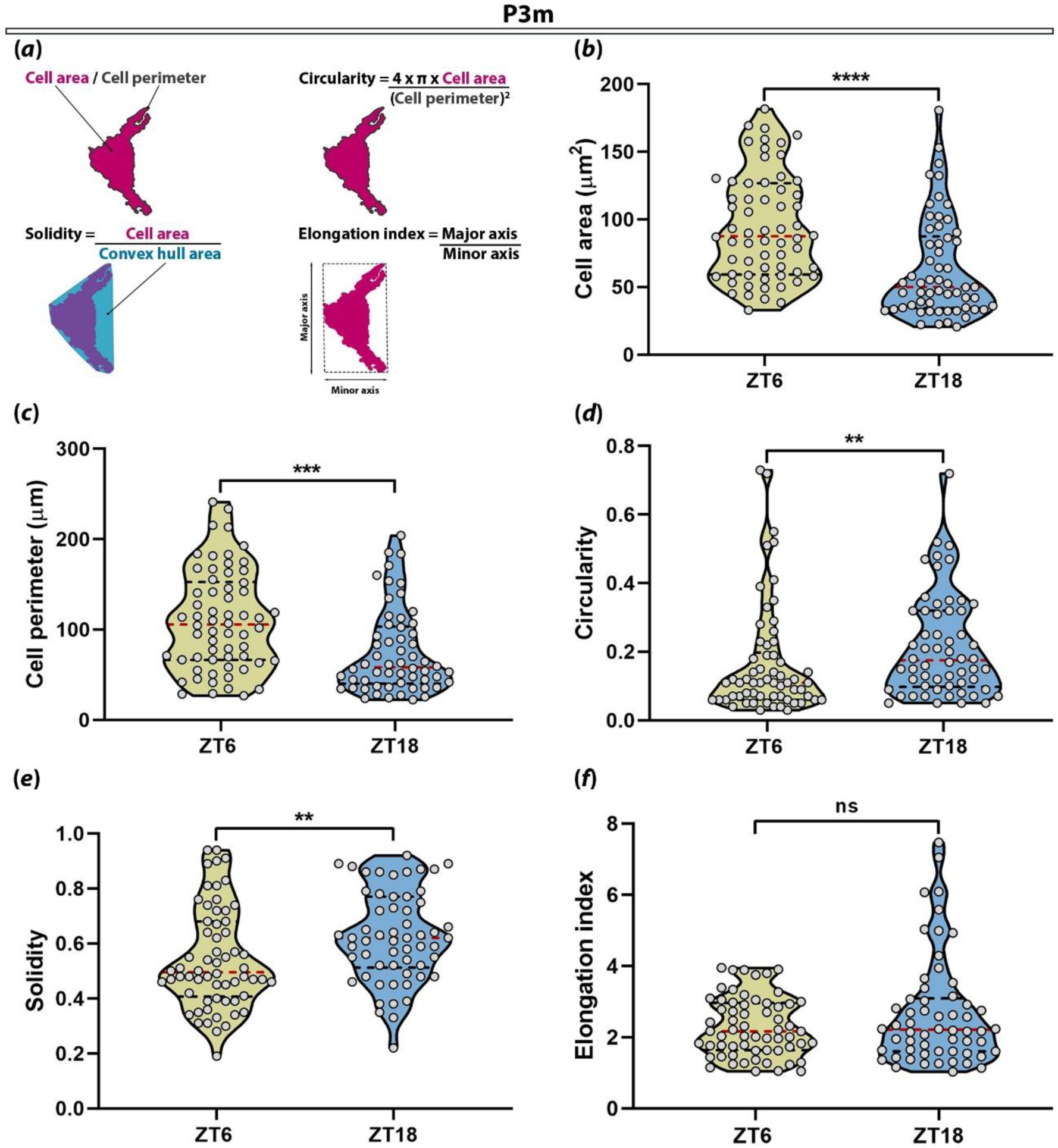
Morphological characterization of IBA1^+^ cells. **(a)** Illustrations and formulas of the morphological parameters calculated in individualized IBA1^+^ cells. **(b)** Cell area (Mann-Whitney U test; ****:p<0.0001). **(c)** Cell perimeter (Mann-Whitney U test; ***:p<0.0002). **(d)** Circularity (Mann-Whitney U test; **:p<0.0060). **(e)** Solidity (Mann-Whitney U test; **:p<0.0065). **(f)** Elongation index (Mann-Whitney U test; p=0.5692). Data were expressed as mean ± SEM. The red dashed line in violin plots indicates the median, while the black dashed lines indicate the quartiles. ns: not significant. P3m: 3-month-old.

### 3.3 Content of lysosomes in pineal IBA1^+^ cells

As the brain’s primary phagocytes, microglia execute a range of biological functions determined by their phagocytic capacity, including synaptic pruning, clearance of apoptotic cells, and immune removal of aberrant elements and pathogens, primarily via lysosome-mediated pathways (Arcuri et al., 2017; Solé-Domènech et al., 2016). Additionally, it has been demonstrated that microglia exhibit rhythmic phagocytic activity, as seen in the elimination of synapses during sleep (Choudhury et al., 2020; Griffin et al., 2020). To assess the lysosomal content in pineal IBA1^+^ cells within the P3m PG at ZT6 and ZT18, we co-stained gland sections for IBA1 and the myeloid-specific lysosomal marker cluster of differentiation 68 (CD68) (Figure 3). The immunostaining revealed that the majority of IBA1^+^ cells were not immunoreactive for CD68 (IBA1^+^ CD68^-^ cells) at both ZTs. On the other hand, IBA1^+^ cells expressing CD68 (IBA1^+^ CD68^+^ cells) showed a cytoplasmic granular pattern. These cells displayed irregular granules clustered or scattered throughout the cytoplasm, particularly in the perinuclear region or within the branches (Figure 3a^1^-a^4^ and 3b^1^-b^4^).

**Figure 3.**
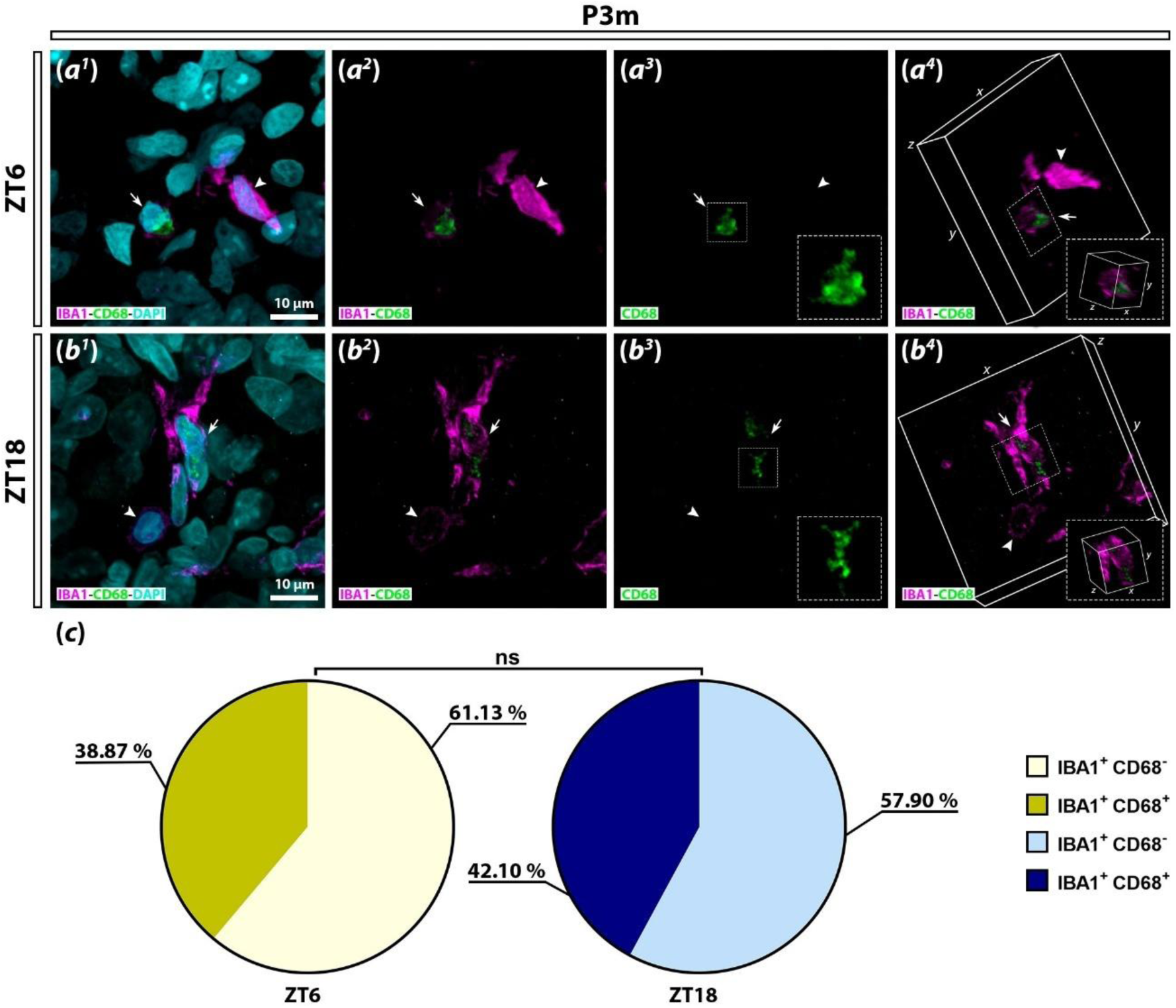
Expression of CD68 in IBA1^+^ cells. **(a-b)** Representative confocal micrographs of gland sections immunolabeled for IBA1 (magenta) and CD68 (green). DAPI (cyan) was used as nuclear marker. (a^1^-a^3^ and b^1^-b^3^) Images taken at 60X with 3.8x digital zoom. Amplified images of the inset boxes are shown in the bottom right corners. (a^4^ and b^4^) Volumetric views of IBA1^+^ cells shown in a^2^ and b^2^, respectively. Amplified images of the inset boxes are shown in the bottom right corners. IBA1^+^ CD68^-^ cells are indicated by white arrowheads and IBA1^+^ CD68^+^ cells by white arrows. **(c)** Percentages of IBA1^+^ cells without immunoreactivity for CD68 (IBA1^+^ CD68^-^; Student’s t test; p=0.8276) and with immunoreactivity for CD68 (IBA1^+^ CD68^+^; Student’s t test; p=0.8276). Percentages shown in the pie charts are the means. Data were expressed as mean ± SEM. ns: not significant. P3m: 3-month-old.

The quantification showed that IBA1^+^ CD68^-^ cells (61.13 ± 8.06 % and 57.90 ± 12.55 % for PGs collected at ZT6 and ZT18, respectively; p=0.8276) outnumbered IBA1^+^ CD68^+^ cells (38.87 ± 18.02 % and 42.10 ± 25.11 %; p=0.8276). Nevertheless, no significant differences were observed in the percentages of IBA1^+^ CD68^-^ cells and IBA1^+^ CD68^+^ cells between both ZTs (Figure 3c).

### 3.4 Autophagy within the pineal organ

The removal and recycling of damaged organelles and misfolded proteins through autophagy are crucial for maintaining normal cellular function and energy balance (Kim & Lee, 2014; Mizushima & Komatsu, 2011). During this process, cytoplasmic materials are carried into the lysosomes for degradation (Jülg et al., 2021; Yim & Mizushima, 2020). As a fundamental mechanism for cellular homeostasis, autophagy is regulated by various factors, including the CTS (Wang et al., 2020). We investigated the influence of the L:D cycle on autophagy within the P3m PG by immunolabeling tissue sections against the autophagic marker microtubule-associated protein 1A/1B light chain 3B (LC3B) (Figure 4). The staining revealed a widespread immunoreactivity for LC3B, displaying a cloudy pattern at both ZTs. Pinealocytes and IBA1^+^ cells showed a higher expression of this marker than IBA1^-^ interstitial cells. However, LC3B expression pattern in IBA1^+^ cells was relatively similar for both ZT6 and ZT18 (Figure 4b^1^-b^4^ and 4c^1^-c^4^).

**Figure 4.**
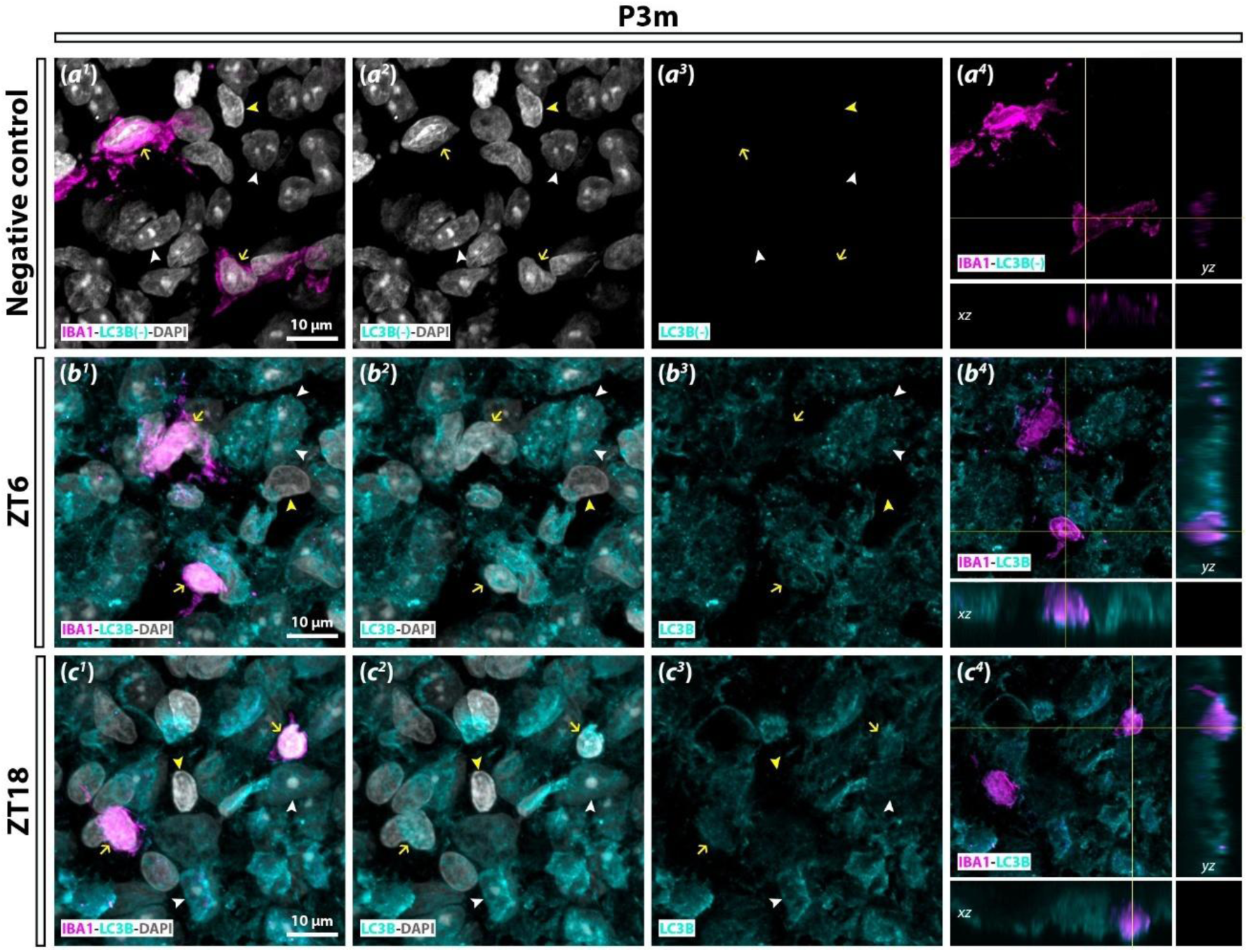
Expression of LC3B within pineal tissue. (**a-c**) Immunostaining of gland sections for IBA1 (magenta) and LC3B (cyan). DAPI (grey) was used as nuclear marker. (a^1^-c^4^) Images at 60X with 3.8x digital zoom. (a^4^, b^4^, and c^4^) Orthogonal views (xz and yz) of single optical planes (1 μm-thick). The orthogonal views correspond to the planes of the intersection points of the yellow lines in each image. (a^1^-a^4^) Negative control of a gland section incubated without the anti-LC3B antibody. IBA1^+^ cells are marked by yellow arrows, IBA1^-^ interstitial cells by yellow arrowheads, and pinealocytes by white arrowheads. P3m: 3-month-old.

### 3.5 Phagocytosis regulation in pineal IBA1^+^ cells

The precise regulation of microglial phagocytosis is indispensable to prevent uncontrolled elimination of cellular elements, which could potentially lead to neurodegeneration (Butler et al., 2021). Activation of modulatory receptors, such as signal regulatory protein alpha (SIRP alpha), initiates signaling pathways in microglial cells that suppress their phagocytic capacity (Feng et al., 2023; Zhang et al., 2015). Notably, SIRP alpha transcript levels have been shown to be upregulated at daytime compared to nighttime in α-microglia from the rat PG (Mays et al., 2018). This study, to the best of our knowledge, provides the first immunolabeling evidence of SIRP alpha expression in pineal cell populations, including IBA1^+^ cells, at both ZTs (Figure 5). SIRP alpha exhibited a finely granular pattern distributed throughout the cytoplasm and periphery of the cells within the P3m PG (Figure 5b^1^-b^4^ and 5c^1^-c^4^). In IBA1^+^ cells, the distribution of SIRP alpha appeared polarized towards specific sites, likely in plasma membrane domains, and specially at daytime (Figure 5b^1^-b^4^). The quantification of the mean fluorescence intensity (286.6 ± 17.5 AU/IBA1^+^ cell and 208.1 ± 11.0 AU/IBA1^+^ cell for glands collected at ZT6 and ZT18, respectively; p=0.0005) confirmed a significant decrease in the signal of SIRP alpha in IBA1^+^ cells at ZT18 compared to ZT6 (Figure 5d).

**Figure 5.**
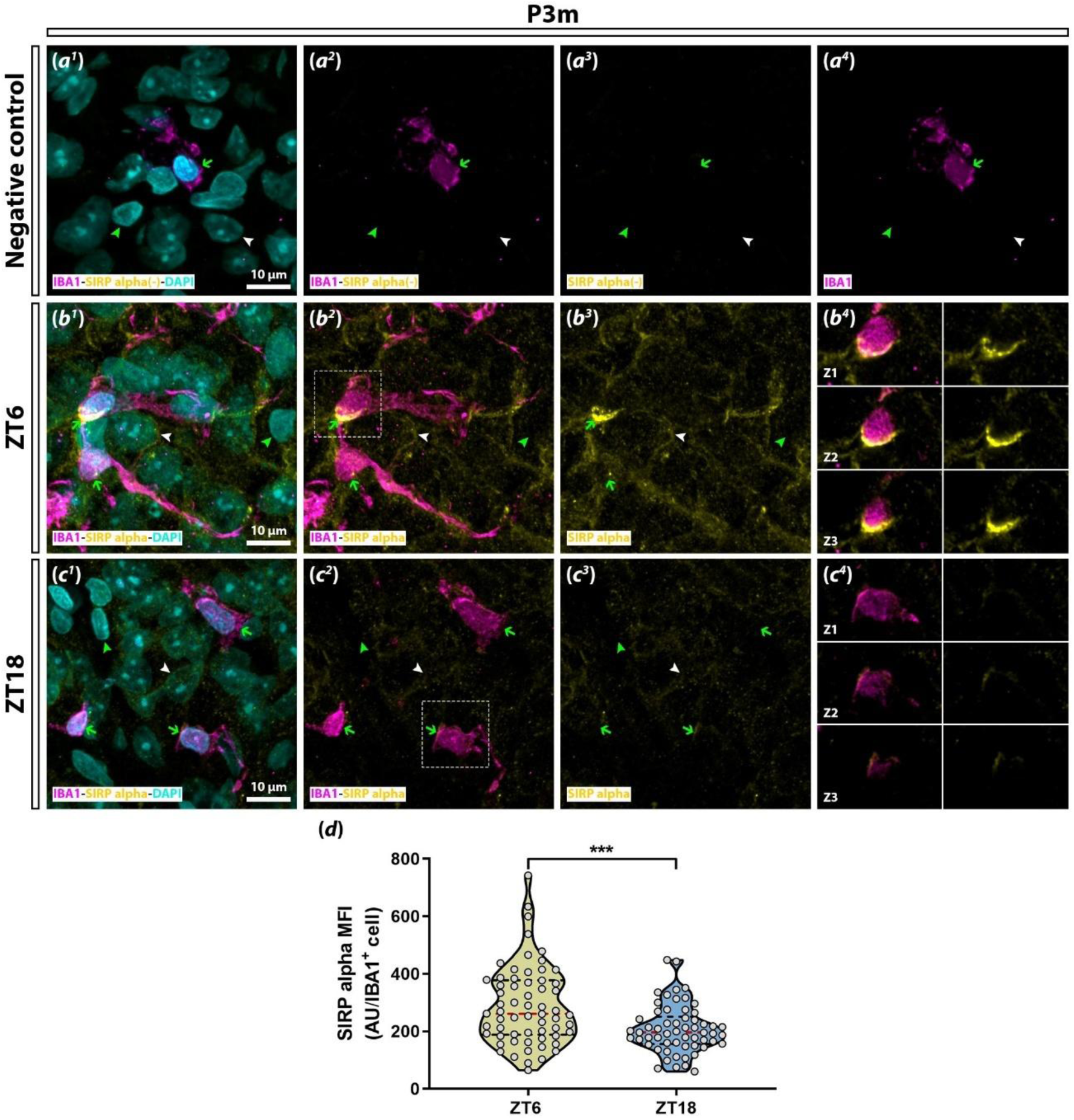
Expression of SIRP alpha in IBA1^+^ cells. **(a-c)** Representative images of gland sections immunolabeled for IBA1 (magenta) and SIRP alpha (yellow). DAPI (cyan) was used as nuclear marker. (a^1^-c^4^) 60X images with 3.8x digital zoom. (b^4^ and c^4^) 1 μm-thick images corresponding to successive optical planes (Z1, Z2, and Z3) taken from the inset boxes in b^2^ and c^2^, respectively. (a^1^-a^4^) Negative control of a gland section incubated without the anti-SIRP alpha antibody. IBA1^+^ cells are indicated by green arrows, IBA1^-^ interstitial cells by green arrowheads, and pinealocytes by white arrowheads. **(d)** Mean fluorescence intensity (MFI) of SIRP alpha signal in IBA1^+^ cells (Mann-Whitney U test; ***:p=0.0005). Data were expressed as mean ± SEM. The red dashed line in violin plots indicates the median, while the black dashed lines indicate the quartiles. AU: arbitrary units. P3m: 3-month-old.

### 3.6 Interactions between IBA1^+^ cells and PAX6^+^ cells within the pineal organ

The developing rodent PG is primarily composed of paired box 6 (PAX6)/vimentin-positive neuroepithelial cells (Borregón et al., 1993; Ibañez Rodriguez et al., 2016; Rath, 2024). The homeobox transcription factor PAX6 is essential for gland ontogeny, with its expression gradually decreasing during development (Rath et al., 2013). However, PAX6^+^ cells persist as a distinguishable population in the adult organ (Ibañez Rodriguez et al., 2016, 2018). Our immunostaining revealed the presence of numerous PAX6^+^ cells sparsely distributed within the P3m PG at both ZTs, with no apparent influence of the L:D cycle on their spatial distribution (Figure 6a^1^-a^3^ and 6c^1^-c^3^).

**Figure 6.**
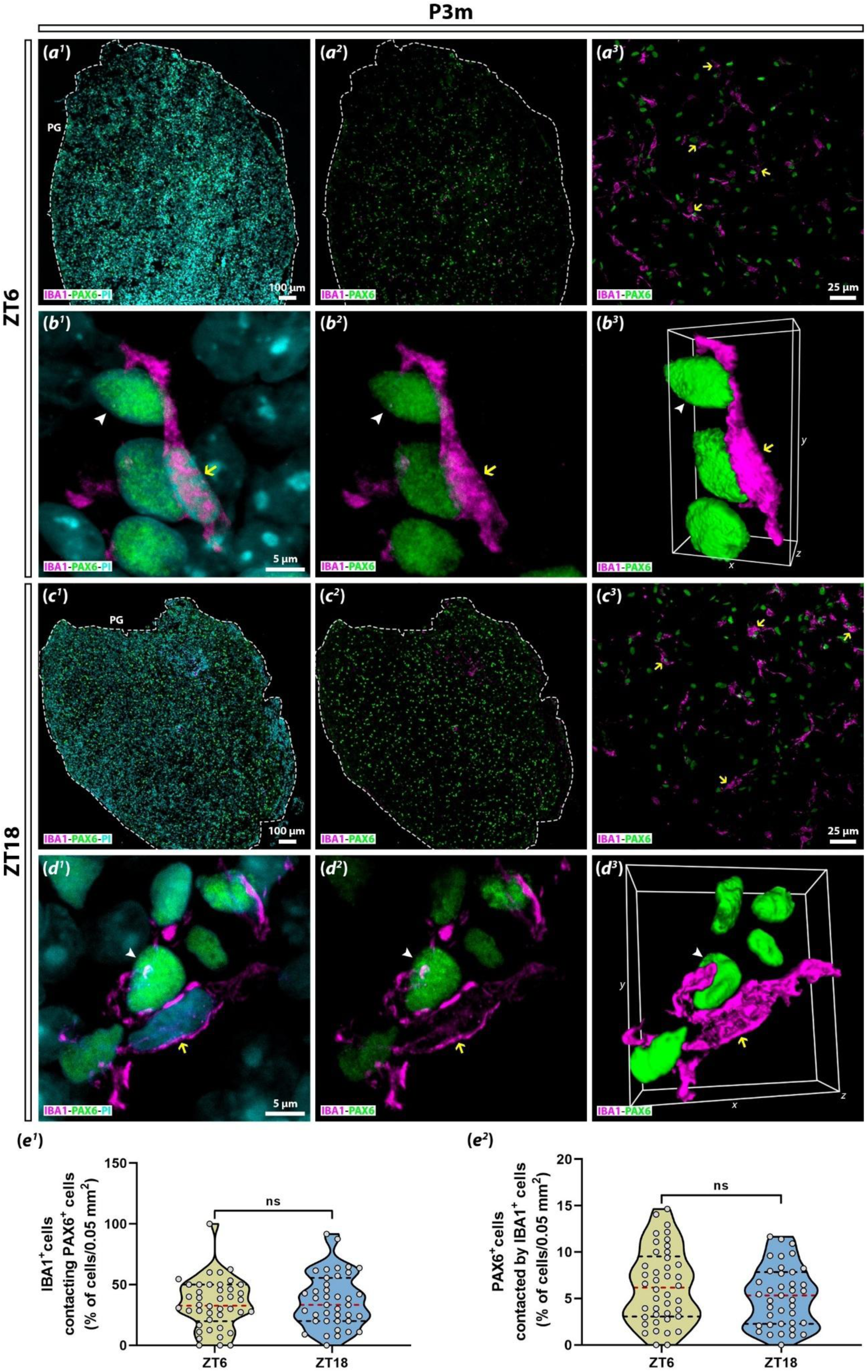
Contact events between IBA1^+^ cells and PAX6^+^ cells. **(a-d)** Representative images of gland sections immunolabeled for IBA1 (magenta) and PAX6 (green). PI (propidium iodide; cyan) was used as nuclear marker. (a^1^, a^2^, and c^1^, c^2^) Images taken at 10X. The perimeter of each gland is delimited by a white dashed line. (a^3^ and c^3^) Images at 40X. IBA1^+^ cells contacting PAX6^+^ cells are denoted by yellow arrows. (b^1^, b^2^, and d^1^, d^2^) Images at 100X with 3.8x digital zoom. IBA1^+^ cells contacting PAX6^+^ cells are indicated by yellow arrows and PAX6^+^ cells by white arrowheads. (b^3^ and d^3^) Three dimensional reconstructions of the cells shown in b^2^ and d^2^, respectively. **(e)** Quantification of contact events between IBA1^+^ cells and PAX6^+^ cells. (e^1^) Percentage of IBA1^+^ cells contacting PAX6^+^ cells (Student’s t test; p=0.5478). (e^2^) Percentage of PAX6^+^ cells contacted by IBA1^+^ cells (Student’s t test; p=0.1757). Data were expressed as mean ± SEM. The red dashed line in violin plots indicates the median, while the black dashed lines indicate the quartiles. ns: not significant. PG: pineal gland. P3m: 3-month-old.

Microglia establish intimate contacts and eventually phagocytize PAX6^+^ cells, from the embryonic to the mature rat PG, under physiological conditions (Ibañez Rodriguez et al., 2016). To engulf neighboring cells, these pineal phagocytes frequently extend one or two thick main processes from the cell body, displaying mono-or bipolar shapes with scarce secondary or tertiary arborization. Considering that microglia and macrophages exhibit rhythmic behaviors in their interactions for tasks such as trimming synaptic elements and removing abnormal materials in other brain regions (Choudhury et al., 2020; Takayama et al., 2016), we analyzed the contact events between PAX6^+^ cells and IBA1^+^ cells in the P3m PG at ZT6 and ZT18, by using immunofluorescence. At both ZTs, we observed that IBA1^+^ cells extended one or two branches to surround nearby PAX6^+^ cells (Figure 6b^1^-b^3^ and 6d^1^-d^3^). Furthermore, a few PAX6^+^ cells were observed to be engulfed by IBA1^+^ cells.

We found that approximately one third of the IBA1^+^ cell population interacted with PAX6^+^ cells at both ZTs (34.09 ± 3.14 %/0.05 mm^2^ and 36.93 ± 3.53 %/0.05 mm^2^ for samples collected at ZT6 and ZT18, respectively; p=0.5478) (Figure 6e^1^). Additionally, low percentages of PAX6^+^ cells were contacted by IBA1^+^ cells (6.35 ± 0.61 %/0.05 mm^2^ and 5.24 ± 0.52 %/0.05 mm^2^; p=0.1757) (Figure 6e^2^). Nevertheless, our statistical analysis did not show significant differences in the cell-cell contacts between the two ZTs studied.

### 3.7 Effects of aging on rhythmic parameters of pineal IBA1^+^ cells

Aging dampens the rhythmic activity of glial cells, including microglia, potentially contributing to neuronal environment deterioration and the progression of age-related diseases (Chen et al., 2023). Therefore, we evaluated whether daily variations in microglial features previously described in the young adult PG were altered in the aged gland. The analysis revealed a significant reduction in cell area (78.58 ± 4.21 μm^2^ and 61.80 ± 4.24 μm^2^; p=0.0012) and cell perimeter (65.57 ± 4.78 μm and 57.47 ± 4.95 μm; p=0.0417) of IBA1^+^ cells from PGs of P18m rats at ZT18 compared to ZT6 (Figure 7a and 7b). On the contrary, circularity (0.34 ± 0.02 and 0.37 ± 0.03; p=0.4539), solidity (0.71 ± 0.02 and 0.72 ± 0.02; p=0.65), and elongation index (2.23 ± 0.14 and 2.08 ± 0.09; p=0.7252), did not show any variation between both ZTs in aged PGs (Figure 7c, 7d, and 7e). It is worth noting that cell area and perimeter decreased by 32.92 % and 31.46 %, respectively, from ZT6 to ZT18 in P3m glands (Figure 2b and 2c). In contrast, in P18m glands, cell area and cell perimeter were reduced by 21.35 % and 12.35 %, respectively, from daytime to nighttime (Figure 7a and 7b). Moreover, the cell area and perimeter means were consistently lower in P18m glands compared to P3m glands (Figures 2 and 7). In terms of microglial morphology diversity, IBA1^+^ cells in the aged gland displayed numerous morphotypes, similar to those observed in the young adult gland (Suppl. Figure 2).

**Figure 7.**
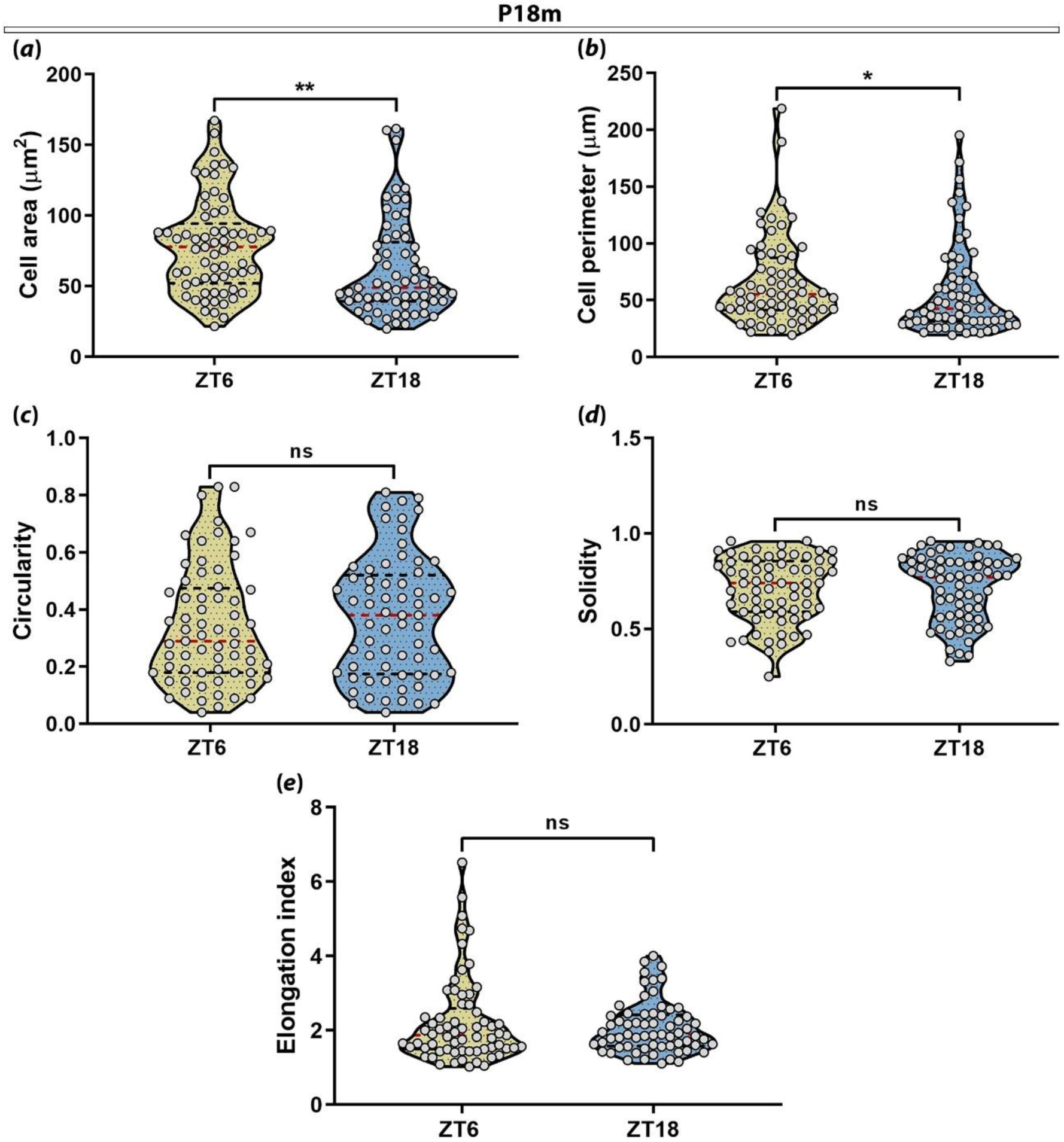
Morphological characterization of IBA1^+^ cells within the P18m PG. (**a**) Cell area (Mann-Whitney U test; **:p=0.0012). (**b**) Cell perimeter (Mann-Whitney U test; *:p=0.0417). **(c)** Circularity (Mann-Whitney U test; p=0.4539). (d) Solidity (Mann-Whitney U test; p=0.65). **(e)** Elongation index (Mann-Whitney U test; p=0.7252). Data were expressed as mean ± SEM. The red dashed line in violin plots indicates the median, while the black dashed lines indicate the quartiles. ns: not significant. P18m: 18-month-old.

Aging also drives dysregulation in circadian and immune response-related genes, facilitating transitions of microglial cells into hyperreactive and pro-inflammatory phenotypes (Costa et al., 2021; Pan et al., 2020). Given that we demonstrated a rhythmic expression of SIRP alpha in IBA1^+^ cells from P3m glands, we subsequently stained sections from P18m PGs to determine whether this rhythmicity is affected by aging (Figure 8). SIRP alpha signal was lesser at ZT18, as compared to ZT6 in aged PGs (Figure 8b^1^-b^4^ and 8c^1^-c^4^). IBA1^+^ cells exhibited a diffuse cytoplasmic pattern, with some areas of greater intensity at the cell periphery. Interestingly, the fluorescent signal was substantially attenuated in the aged samples compared to the young adult samples at the corresponding ZTs (Figures 5 and 8).

**Figure 8.**
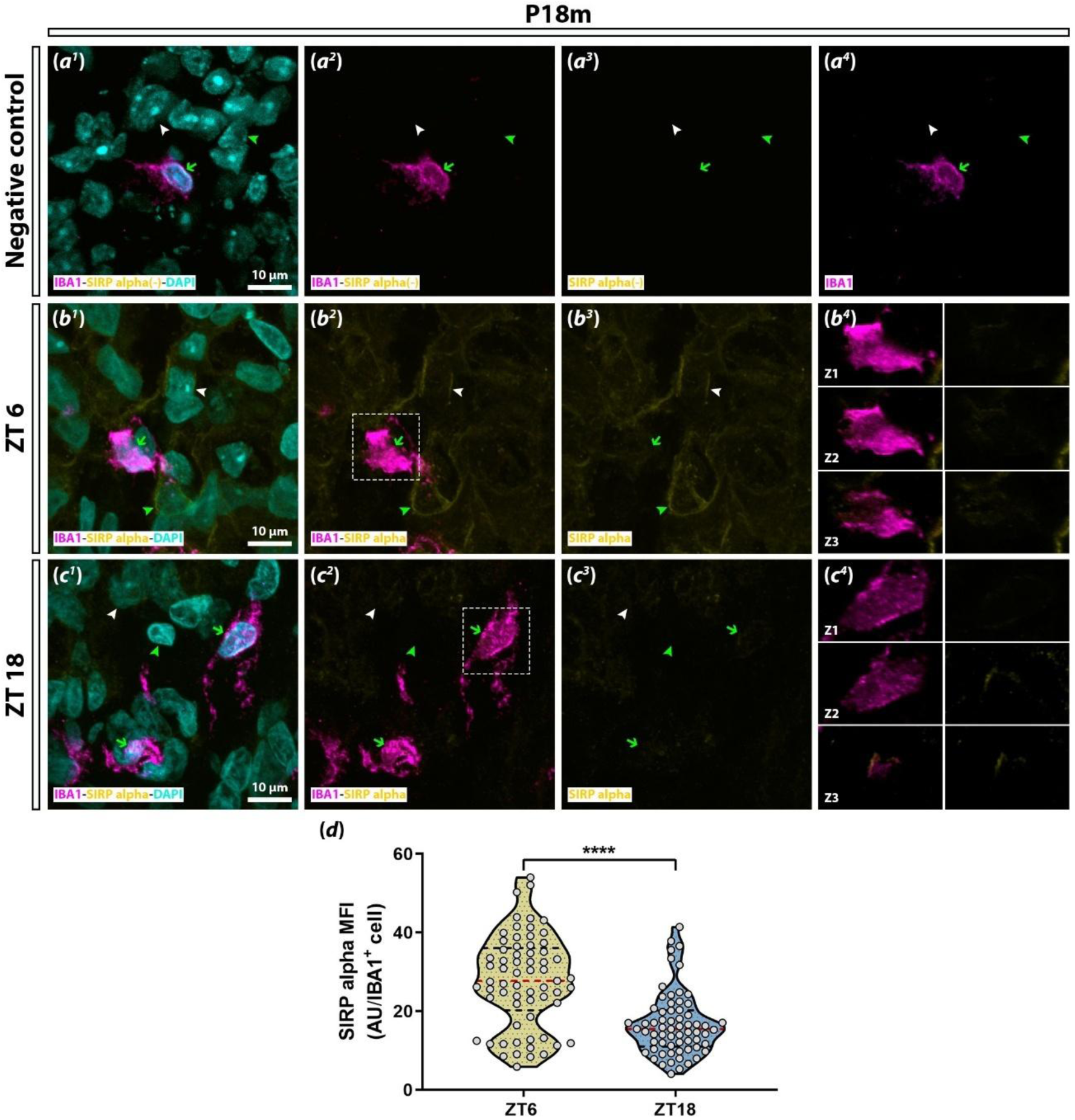
Expression of SIRP alpha in IBA1^+^ cells within the P18m PG. **(a-c)** Representative images of gland sections immunolabeled for IBA1 (magenta) and SIRP alpha (yellow). DAPI (cyan) was used as nuclear marker. (a^1^-c^4^) 60X images with 3.8x digital zoom. (b^4^ and c^4^) 1 μm-thick images corresponding to successive optical planes (Z1, Z2, and Z3) taken from white dashed line areas in b^2^ and c^2^, respectively. (a^1^-a^4^) Negative control of a gland section incubated without the anti-SIRP alpha antibody. IBA1^+^ cells are indicated by green arrows, IBA1^-^ interstitial cells by green arrowheads, and pinealocytes by white arrowheads. **(d)** Mean fluorescence intensity (MFI) of SIRP alpha signal in IBA1^+^ cells (Mann-Whitney U test; ****:p<0.0001). The red dashed line in violin plots indicates the median, while the black dashed lines indicate the quartiles. AU: arbitrary units. P18m: 18-month-old.

Fluorescence quantification confirmed a daily variation in SIRP alpha signal in aged IBA1^+^ cells, with a significant reduction at midnight (27.74 ± 1.45 AU/IBA1^+^ cell and 16.71 ± 1.00 AU/IBA1^+^ cell; p<0.0001) (Figure 8d). A substantial age-related difference was also confirmed by this analysis. In P3m PGs, SIRP alpha intensity decreased by 27.39 % from ZT6 to ZT18, whereas in P18m glands, the reduction reached 39.76 %. The SIRP alpha intensity mean in IBA1^+^ cells from aged glands was lower compared to that in young adults (Figures 5d and 8d).

## 4. Discussion

Nowadays, it is widely accepted that microglia are a multifaceted and plastic cell population within the CNS, adapting their morphology and physiology in response to multiple factors, both local and external (Augusto-Oliveira et al., 2022; Paolicelli et al., 2022; Shemer et al., 2015). The sensitivity of microglial cells to detect environmental changes is significantly heightened in brain regions where there is a limited regulation of molecule exchange between the bloodstream and nervous tissue. These brain areas, known as circumventricular organs (CVOs), are neuroepithelial structures located close to the third and fourth ventricles, and they lack a complete blood brain barrier because their capillaries are fenestrated (Kiecker, 2018; Miyata, 2015; Szathmari et al., 2015). Due to these anatomical characteristics, microglia within the CVOs, that include the PG, operate as highly specialized sensors able to detect and react appropriately to diverse molecular signals (Ibañez Rodriguez et al., 2018; Ott et al., 2010; Takagi et al., 2019; Vollmer et al., 2016; Wuchert et al., 2008).

The CTS exerts notable regulation on microglia behavior, by providing a temporal order to their functions. In this way, microglia efficiently carry out immune responses against harmful agents and perform basal tasks under homeostatic conditions (Brécier et al., 2023; Jiao et al., 2024). In this study, we characterized IBA1^+^ microglial cells within the young adult and aged rat PG at two points of the L:D cycle by applying both qualitative and quantitative analyses of confocal microscopy data.

The spatial distribution of microglia is heterogeneous across the CNS (Stratoulias et al., 2019; Tan et al., 2020). For instance, there is evidence that microglial cells are more densely distributed in anterior and medial parts of the mammalian brain, such as the olfactory bulb, cortex, hippocampus, basal ganglia, and substantia nigra, as compared to posterior regions like the cerebellum, brainstem, and spinal cord (Lawson et al., 1990; van Weering et al., 2023).

In the young adult PG, our counting showed that IBA1^+^ cells were randomly dispersed throughout the pineal tissue, either as individual cells or as cell clusters, with no significant differences between ZT6 and ZT18. Similarly, it was demonstrated that the total number of IBA1^+^ cells did not vary within the cortex and hippocampus of wild type mice between the light and night phases (Griffin et al., 2019; Hayashi et al., 2013). Nevertheless, the density of microglial cells in the murine arcuate nucleus significantly raised at ZT16 in contrast to ZT4, in a 24 h L:D cycle, in both normal and hypercaloric conditions. The increase in microglia numbers may enhance their interactions with arcuate pro-opiomelanocortin neurons during the feeding time, prompting this neuronal population into highly energy-demanding states (Yi et al., 2017). Herein, we hypothesize that changes in microglia number may not be required to modulate the rhythmic activity of pinealocytes under homeostatic conditions. Unlike pro-opiomelanocortin neurons, pinealocytes would optimally self-regulate with a stable number of dynamic microglial cells under physiological conditions.

Migration, recruitment, and formation of cell aggregates constitute a typical response of microglia to injuries (Clark, 1974; Marella & Chabry, 2004; Paasila et al., 2020; van Horssen et al., 2012). Therefore, the cause of microglial cluster formation within the PG under homeostatic conditions has yet to be clarified. In the present study, we did not find differences in the number of IBA1^+^ cell clusters between ZT6 and ZT18 in the P3m PG.

Daily changes in microglial morphology have been documented in different brain regions. For instance, murine cortical microglia exhibited greater total length of processes and number of branch points during the night phase (Hayashi et al., 2013; Takayama et al., 2016). It has been postulated that microglia, activated by neuronal ATP via P2Y_12_ receptors, become hyper-ramified during wakefulness (nighttime), thereby enhancing interactions with cortical neurons. These microglia secrete the proteolytic enzyme cathepsin S near dendritic spines, dissolving the perineural nets and regulating synaptic strength at the onset of sleep (daytime), which is essential for the acquisition of new information (Nakanishi et al., 2021). In other areas related to the initiation and propagation of slow waves—rhythmic oscillations produced during sleep—such as the somatosensory cortex, dorsal hippocampus, and basal forebrain, microglia displayed reduced cell volume and ramification at daytime (Steffens et al., 2023). The authors proposed that local vigilance states of neurons and slow waves, both occurring in the light phase, modulate these microglial de-ramified shapes (Steffens et al., 2023). These results show that neuronal activity could modulate microglial morphology. Unlike the aforementioned, IBA1^+^ cells in the young adult PG exhibited significantly smaller cell area and perimeter at ZT18 compared to ZT6. Moreover, the cells presented greater values of circularity and solidity at ZT18. These data suggest that the daily fluctuations in microglial shape may be influenced by factors with immunomodulatory activity within the PG. *In vitro* and *in vivo* studies conducted in different rodent brain regions have demonstrated that administrated MEL triggers morphological shifts in microglial cells, leading to amoeboid or rounded shapes (Kaur & Ling, 1999; Merlo et al., 2023). In addition, MEL also exerts suppressive effects on inflammatory responses of microglial cells, stimulating polarization into anti-inflammatory phenotypes (Azedi et al., 2019; Gao et al., 2024; Wen et al., 2016; Yao et al., 2019). Moreover, microglial cells in the murine visual cortex exhibited more complex arborization when animals were anesthetized, as compared to when they were awake (nighttime), a phase marked by high norepinephrine levels. Interestingly, pharmacological stimulation of β_2_-adrenergic receptors mimics these effects (Stowell et al., 2020). Considering that norepinephrine levels begin to rise after the dark onset within the rat PG and MEL levels peak around midnight (ZT18; the time of sacrifice of our animals) (Borjigin et al., 2012; Drijfhout et al., 1996; Maronde et al., 1999), these hormonal modulators may induce pineal microglial cells to adopt non-inflammatory reactive phenotypes. Such phenotypes are likely oriented towards tissue repair and homeostatic tasks, accompanied by rounded morphologies. Even though phagocytes within the CVOs are scarcely ramified (Muñoz, 2022), these small, amoeboid-like microglia could easily navigate through the pineal tissue to carry out beneficial functions at night.

Microglia are specialized brain phagocytes able to internalize and degrade harmful or unnecessary elements (Cherry et al., 2015; Cockram et al., 2019; Sierra et al., 2010). In the rat PG, microglia show detectable levels of CD68, a well-established marker of phagocytosis in myeloid cells, which are evident throughout embryonic to postnatal stages. This phagocytic capacity of microglial cells is directly correlated with the progressive decrease in the density of PAX6^+^ precursor cells, suggesting that phagocytes play a regulatory role in modulating precursor cell populations within the PG (Ibañez Rodriguez et al., 2016). Herein, we detected that the majority of IBA1^+^ cells were not reactive for CD68 (IBA1^+^ CD68^-^ cells) within the young adult PG. However, the IBA1^+^ cells positive for this marker (IBA1^+^ CD68^+^ cells) showed CD68^+^ clumps of different sizes. No significant differences were found in the percentages of these two microglial phenotypes between both ZTs analyzed. Clearance activity of microglial cells also exerts daily oscillations. It has been demonstrated that microglia from murine prefrontal cortex eliminate synapses, as evidenced by the presence of CD68^+^ phagosomes containing the synaptic protein synaptophysin. This process was found to be more pronounced at ZT0 than at ZT12 in a 24 h L:D cycle. The authors claimed that microglia perform synaptic pruning of unnecessary or inactive synapses at the beginning of the light phase (sleep phase) (Choudhury et al., 2020). In the young adult rat PG, the absence of daily variations in the quantity of CD68-reactive IBA1^+^ cell phenotypes at the two ZTs analyzed may indicate that pineal phagocytes do not execute waste removal in a temporally regulated manner. However, we acknowledge that further analysis, covering additional time points across the L:D cycle, is required to more accurately assess the rhythmicity of the pineal microglia’s phagocytic capacity.

Microglial cells within the CVOs constantly patrol their microenvironment, maintaining reactive states, and showing intense expression of CD68, even under physiological conditions (Takagi et al., 2019). Furthermore, in pathological contexts, peripheral immune challenges can impact on microglial physiology in the CVOs, triggering robust and rapid activation, suggesting that these regions are the primary sites for the initiation of the microglial immune response within the brain (Gaige et al., 2022; Kloss et al., 2001; Matsushita et al., 2021).

Circadian regulation of autophagic processes within the rat PG may impact on its physiology. Pinealocytes exhibit increased levels of mitochondrial autophagy (mitophagy) during daytime, leading to a reduction in mitochondrial number and a subsequent depletion of ATP (Ge et al., 2021). This reduces cyclic AMP availability, which is crucial for regulating the expression of *Aanat* (arylalkylamine N-acetyltransferase), the gene that encodes AANAT, the penultimate enzyme involved in MEL synthesis (Klein, 2007). The authors hypothesized that lower cyclic AMP levels result in downregulation of *Aanat*, contributing to the suppression of the enzyme production during the day (Ge et al., 2021). In this study, we showed that the signal of the autophagic marker LC3B appeared diffusely distributed in IBA1^+^ cells at ZT6 and ZT18. Unlike pinealocytes, where autophagy seems to be organized in a time-dependent manner to optimize MEL production at night, pineal IBA1^+^ cells appear to sustain a consistent basal autophagic flux throughout both day and night. These findings suggest that pineal microglial cells may remain in reactive states associated with material recycling and clearance across the L:D cycle.

Microglial phagocytosis is regulated by receptor-ligand interactions such as the CD47-SIRP alpha axis. When the microglial SIRP alpha receptor binds to its ligand CD47, expressed in several cell populations, phagocytosis is repressed. It has been demonstrated that CD47-SIRP alpha signaling is essential to prevent an excessive synapse pruning in the developing brain (Jiang et al., 2022; Lehrman et al., 2018). We found herein that SIRP alpha signal was widespread across the young adult PG at both ZT6 and ZT18, but with lower levels at midnight. In IBA1^+^ cells, SIRP alpha expression was mostly cytoplasmic, although also in the cell periphery, presumably in membrane domains, as shown before in cultured primary microglial cells (Gitik et al., 2011). The fluorescence intensity measurement confirmed a statistically significant decrease in SIRP signal at ZT18 in IBA1^+^ cells from the young adult PG. It is important to note that it has been speculated that tissue dissociation, commonly used to analyze the transcriptomic profiles of specific cell populations, may disrupt or modify the oscillatory expression of certain genes. In this study, we provided *ex vivo* evidence that IBA1^+^ cells showed a differential daily expression of SIRP alpha, through immunofluorescence examination. Our result aligns with prior findings at transcriptomic level in microglial cells, which identified *Sirpa* as one of the few rhythmic transcripts downregulated at night in α-microglia within the rat PG (Mays et al., 2018). We propose that the oscillatory expression of SIRP alpha in phagocytes may indicate a protective mechanism within the pineal tissue. It has been demonstrated that bacterial lipopolysaccharide administration on rat PG cultures induces TNF alpha release by microglial cells. This inflammatory mediator might be involved in a signaling cascade within pinealocytes, where it inhibits AANAT expression and consequently MEL synthesis (da Silveira Cruz-Machado et al., 2012). Thus, the attenuated expression of SIRP alpha in IBA1^+^ cells in the dark phase under physiological conditions may enable these phagocytes to mount an effective and rapid immune response against pathogenic microorganisms that could impair MEL production.

The transcription factor PAX6 has been identified within the subcommissural organ, subventricular zone, olfactory bulb, cerebellum, and PG of adult rodents (Corales et al., 2022; Duan et al., 2013; Ibañez Rodriguez et al., 2016; Rath, 2024). PAX6 is expressed predominantly in mature astrocytes and progenitor-like cells in the adult brain (Corales et al., 2022; Mays et al., 2018). Transcriptomic analysis of the rat PG revealed the presence of three astrocyte subtypes, referred to as α-, β-, and γ-astrocytes. Interestingly, *Pax6* showed predominant expression in α– and γ-astrocytes (Mays et al., 2018). These findings imply that IBA1^+^ cells may specifically interact with these astrocytic subpopulations within the young adult PG. In our study, we observed that PAX6^+^ cells were spread throughout the young adult PG at both ZT6 and ZT18. While in the developing rat PG, microglial cells engulf and eliminate PAX6^+^ cells (Ibañez Rodriguez et al., 2016), we showed that around one-third of IBA1^+^ phagocyte population (approximately 34-37 %) interact with PAX6^+^ cells in the young adult gland without significant differences in the contact events between both ZTs. Furthermore, only a small proportion of PAX6^+^ cells (approximately 5-7 %) was contacted by IBA1^+^ cells. Microglia interact with astrocytes in the adult brain to remove dying cortical neurons in a synchronized and spatiotemporal way (Damisah et al., 2020), and to influence synaptic transmission by modulating astrocyte functionality in the cortex and hippocampus (Du et al., 2022; Takata-Tsuji et al., 2021). Moreover, microglial cells play a crucial role in regulating neuronal fate, balancing neuronal death and neurogenesis. For instance, microglia positively regulate precursor cells, promoting their survival, proliferation, and migration in the retina and subventricular zone in the adult mice (Kuse et al., 2018; Ribeiro Xavier et al., 2015). Based on this evidence, we propose that IBA1^+^ cell dynamics with this distinct PAX6-expressing population, whether astrocytes or precursor cells, in the mature rat PG, may influence both cellular identity and physiology, albeit without exhibiting any apparent rhythmic pattern.

As microglia age, they undergo substantial alterations, including circadian disruptions that vary significantly among CNS regions (Damani et al., 2011; Hart et al., 2012; Pan et al., 2020; Ritzel et al., 2015). In particular, cultured hippocampal microglia isolated from aged rats exhibited a desynchronized pattern of inflammatory cytokine expression, consistently maintaining elevated levels of *Il1β* and *Tnf* across the day, and in contrast to the rhythmic expression observed in microglia from young animals. These aged microglial cells also showed a lack of daily rhythmicity in the expression of *Aif1* (IBA1-encoding gene) and *Cd68*, showing increased expression of these markers at both ZT6 and ZT18, whereas young cells displayed rhythmic variation (Fonken et al., 2016). As we observed in the young adult PG, IBA1^+^ cells within the aged PG showed a significant reduction in cell area and perimeter at ZT18 compared to ZT6. Nevertheless, these daily changes were less pronounced in P18m glands, suggesting a potential loss of rhythmicity in microglial morphology with aging. Furthermore, in aged samples, SIRP alpha expression in IBA1^+^ cells displayed significant decline at ZT18 in contrast to ZT6. While the daily switch in SIRP alpha signal was more pronounced in aged PGs than in young adult glands, the overall lower intensity of this marker in aged IBA1^+^ cells may reflect dysregulated phagocytosis. These findings suggest that although some IBA1^+^ cell parameters do retain their daily rhythmicity in the aged PG, most of the rhythms were found to be dampened or abolished. Consequently, we do not rule out the possibility that IBA1^+^ cells may be undergoing changes that affect their normal functionality and their impact in the MEL rhythm during aging.

## 5. Concluding remarks

Our findings indicate that pineal IBA1^+^ cells are subject to morphological and functional adaptations influenced by the L:D cycle. These dynamic changes are crucial for effectively responding to harmful stimuli and for maintaining baseline tasks under homeostatic conditions within the rat PG. Future research should aim to determine whether the daily variations observed in microglial parameters are under the CTS control. This could be addressed by employing constant darkness conditions or by disrupting the superior cervical ganglia-derived sympathetic innervation of the PG, either by excision of a segment of the afferent sympathetic trunks (decentralization) or by bilateral removal of the ganglia (bilateral ganglionectomy), among other approaches (Avila et al., 2023; Savastano et al., 2010).

## This PDF file includes

Main text

Figures 1 to 8

Supplementary materials will be available in the final peer-reviewed version.

## Acknowledgements

We would like to express our gratitude to Jorge E. Ibañez for his assistance with confocal microscopy and statistical analysis, Julieta Scelta and Adrián Fernández for their help with animal handling, Andrea A. Bischof for designing the graphical abstract, and Raymond Astrue for editing the manuscript. Funded by the National Agency for the Promotion of Research, Technological Development, and Innovation (ANPCYT), Argentina. Grant/Award Number: PICT2017-0499: EMM and PICT2021-0314: EMM.

